# Identification of enamel knot gene signature within the developing mouse molar

**DOI:** 10.1101/2021.06.14.448115

**Authors:** Emma Wentworth Winchester, Justin Cotney

## Abstract

In most mammals, the primary teeth develop *in utero* and the cells capable of contributing to hard surface regeneration are lost before tooth eruption. These cells differentiate through a series of reciprocal induction steps between the epithelium and mesenchyme, initially orchestrated by an epithelial signaling center called the enamel knot. While the factors secreted by this structure are of interest to the dental regeneration and development communities, its small size makes it difficult to isolate for analysis. Here we describe our work to identify the enamel knot from whole E14 molars using publicly available scRNA-seq data. We identified 335 genes differentially expressed in the enamel knot compared to the surrounding tissues, including known enamel knot marker genes. We validated expression of the most highly enriched enamel knot marker genes and identified 42 novel marker genes of the enamel knot which provide excellent targets for future dental regeneration investigations.

## Background

In most mammals, primary teeth develop *in utero* and have completed most of their formation by birth. The hard tooth surfaces are composed of enamel and dentin, laid down by epithelium- and mesenchyme-derived cells. These cells are not maintained after eruption; as such, mammalian teeth cannot be regenerated *in vivo*. Identifying and harnessing the gene programs that specify these cell types could enable treatment of patients with tooth damage or loss. However, due to the time period in human development that these cell types arise and lack of clear *in vitro* models to identify such gene signatures, tooth development must be studied using primary tissues from mouse development.

While rodent incisors continuously erupt and contain regenerative potential, mouse molars are considered to be representative of most mammalian dentition and development. During mammalian molar formation, sequential stages of reciprocal induction are required for proper derivation of enamel- and dentin-producing cells. (Thesleff, 2003) Around day 13.5 of embryonic development (E13.5), the molar epithelium invaginates into the mesenchyme and a signaling center appears at the epithelial-mesenchymal border. This signaling center, the primary enamel knot (EK), is a group of aproliferative epithelial cells which secrete factors to shape the tooth through tightly controlled proliferation and differentiation of the epithelium and mesenchyme (Vaahtokari *et al*., 1996; Thesleff, Keränen and Jernvall, 2001; Matalova *et al*., 2005). Its loss arrests tooth development and prevents growth and differentiation of dental tissues (Dassule *et al*., 2000), illustrating its importance in tooth development and making it of interest to dental regeneration researchers.

Characterization of the EK transcriptome has been limited as its small size makes isolation difficult. In this work, we show that EK profiling is possible through analysis of publicly available scRNAseq data of E14 mouse molars. Given the large number of cells and replicates (n=4; ~35,000 cells), we hypothesized that the EK, present at E14, could be found in this dataset to identify a robust gene expression signature for use in downstream isolation and manipulation experiments. In this thorough reanalysis, we identified a subcluster of epithelial cells we propose to be the EK. Genes specific to and enriched in these cells are strongly associated with odontogenesis, including many known EK genes. We validated the expression of the most highly enriched EK marker genes using publicly available in situ hybridization data from the E14.5 mouse, which confirmed the specificity of the vast majority of the novel EK markers identified by our analysis. These genes and their overall signature provide excellent targets for future tooth development and regeneration related research.

## Results

### Re-Alignment and Quality Control

Previously published raw reads from four E14 mouse molars (Hallikas *et al*., 2021) were aligned to the mouse genome (mm10) with pseudoaligner kallisto, based on improved performance relative to other approaches (Shainer and Stemmer, 2020). A total of 33,886 cells passed quality control filters for number of genes detected, total counts, and percent of reads derived from mitochondrial gene expression (Fig. 1A-C, Supp. Fig. 1, Methods). Individual replicates were merged, reads were jointly normalized, and gene expression values were scaled to standardize expression variance (Ilicic *et al*., 2016). Having combined the data into a uniformly processed set, we sought to identify major cell types of the developing molar. A graph-based clustering approach (Yuan, 2019) revealed 16 clusters representing distinct cell populations within the developing molar (Fig. 1D). Each biological replicate contributed equally to most clusters, with the exception being cluster 10, demonstrating the overall reproducibility of staging, dissection, and processing of original replicates (Fig. 1E-F).

**Figure 1.**
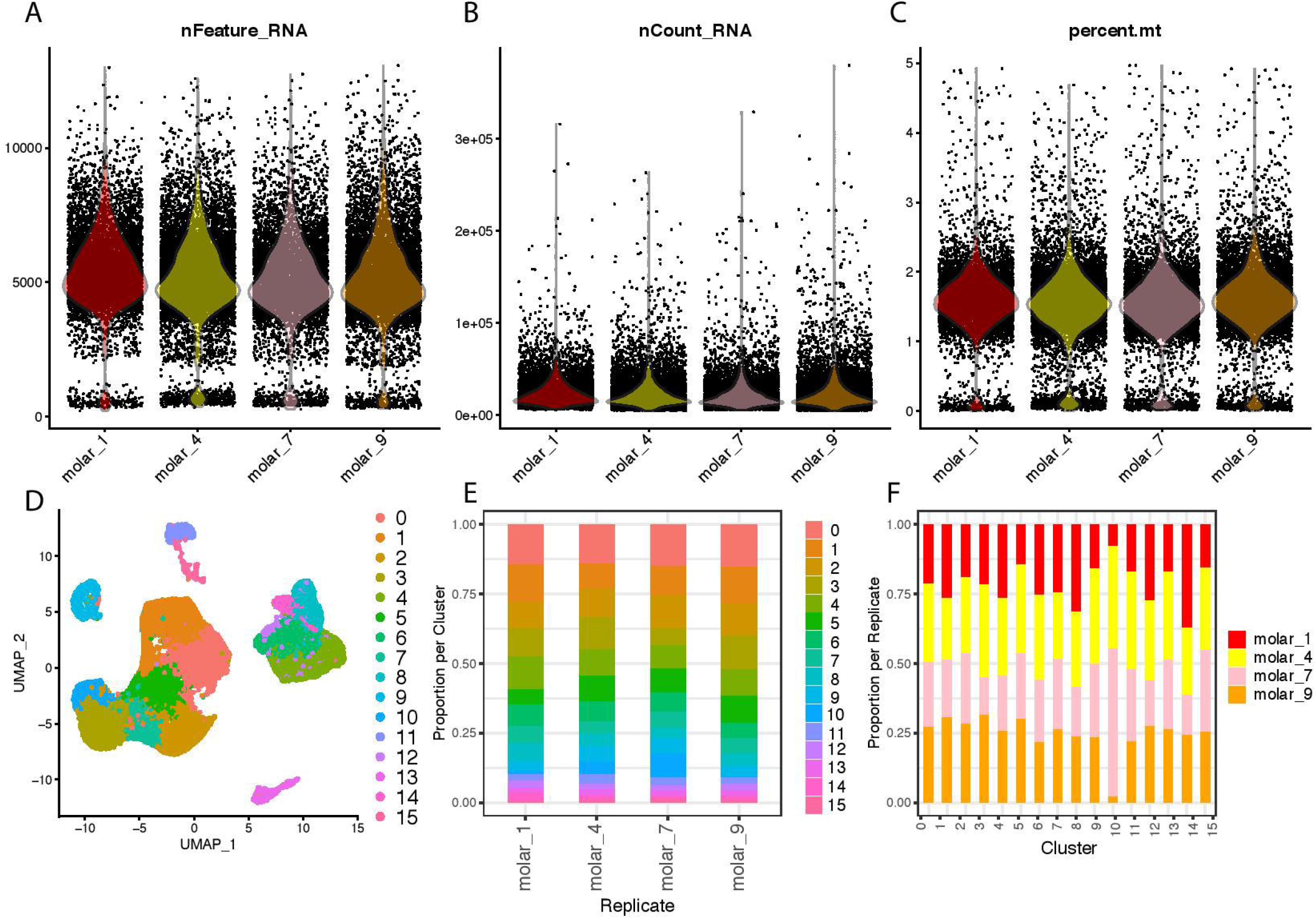
scRNAseq Analysis of E14 Molar Reveals 16 Distinct Cell Populations. **A-C.** Violin plots of per-replicate quality control metrics used to filter cells before analysis. Cells were filtered for number of total aligned reads and percent mitochondrial reads (nFeature > 200 and <5% mitochondrial reads [pct.mt]). **D**. UMAP projection of all 33,886 cells passing quality control filters, with assigned numerical cluster identities from Seurat. **E**. Per-replicate visualization of percent of cells per numerical cluster identity. **F**. Per-cluster visualization of replicate contribution to each numerical cluster identity.

### Classification of cell populations

After identifying 16 distinct cell populations, we sought to classify their biological identities in a systematic, unbiased fashion. We first identified differentially expressed genes that are specific to and enriched within one cluster compared to all others, commonly referred to as marker genes (Andrews *et al*., 2021). We identified 4821 marker genes (average 301/cluster) with 0.5 fold higher average expression in one cluster compared to all others and expressed in at least 25% of cells in that cluster (Methods). We then performed gene ontology (GO) enrichment analysis of these marker genes from each cluster (Supp. Table 1) relative to a background of all genes expressed in this data set (Supp. Table 2) to ascertain biological functions. Using GO annotations associated with biological processes we identified significant GO terms enriched in each cluster (Supp. Table 3). The most significantly and specifically enriched GO term for each cluster is visualized on two dimensional uniform manifold approximation and projection (UMAP) of the single cell data in Fig. 2A.

**Figure 2.**
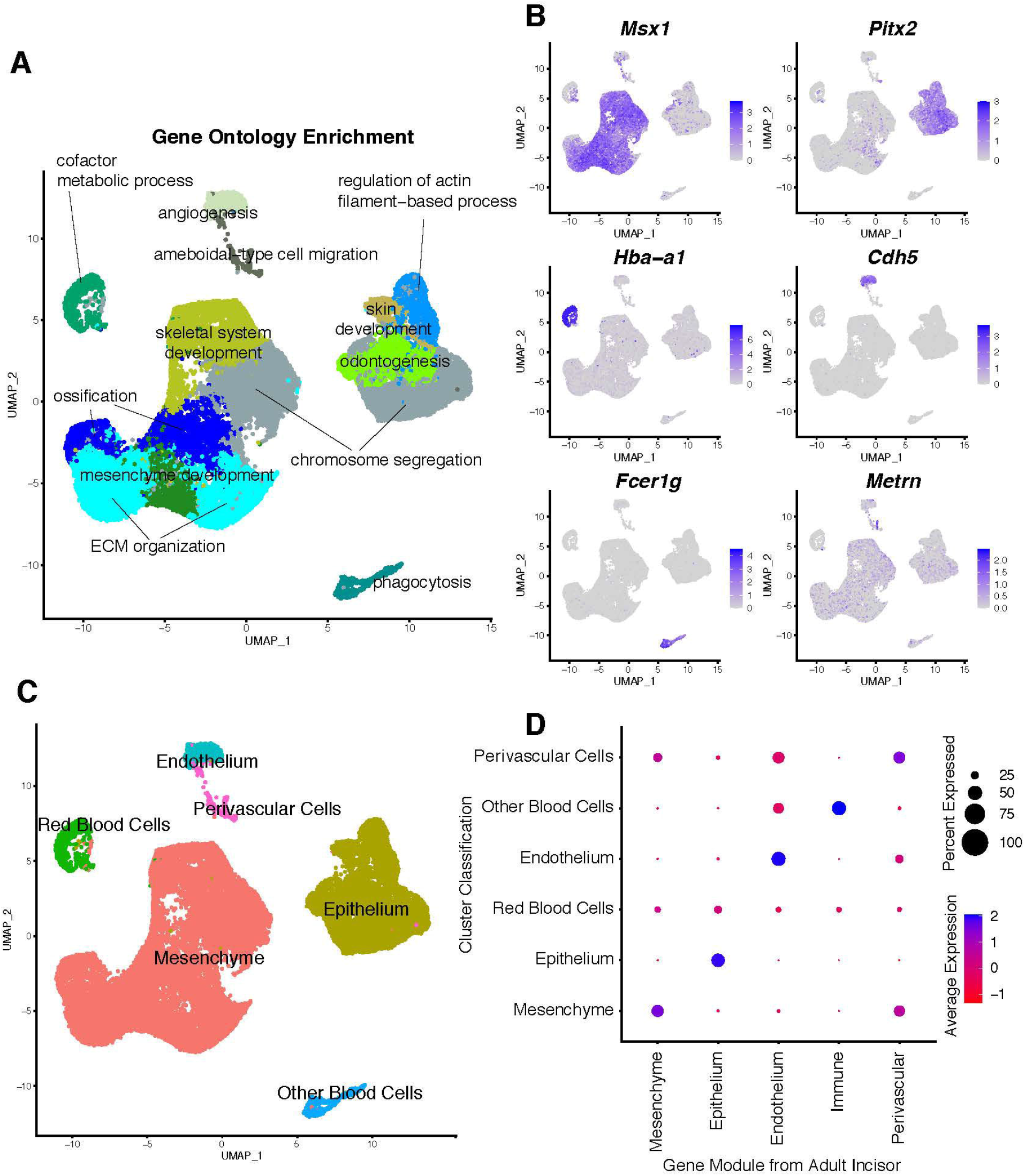
GO-Based Annotation of E14 Molar Cell Types. **A**. UMAP projection overlaid with the most specific highly enriched GO term per cluster. Enriched GO terms were identified using clusterProfiler (Methods). **B**. UMAP projection with overlay of scaled and normalized expression per cell for *Msx1, Pitx2, Hba-a1, Cdh5, Fcer1g*, and *Metrn*, marker genes for mesenchyme, epithelium, red blood cells, endothelial cells, immune cells, and perivascular cells, respectively. **C**. UMAP projection with GO-based cluster annotations. **D.** Dot plot of GO-based cluster classification (y axis) vs gene module from marker genes of published adult mouse incisor cell type (x axis, Krivanek *et al*., 2020 marker genes, average logFC >2). Dot size represents percent of cells in each cluster expressing that gene; blue color represents high enrichment of genes in the module while red represents depletion of genes in the module.

Using this approach, marker genes of 7 tightly linked clusters (0-3,5,7) were enriched for mesenchymal related GO terms, including mesenchymal development and ECM organization (Fig. 2A). To confirm this finding, we examined expression of the dental mesenchymal marker *Msx1* (Jowett *et al*., 1993), which is expressed in all of these populations. (Fig. 2B) Therefore, we classified cells from these clusters as “Mesenchyme” (Fig. 2C). Cluster 10 which was skewed toward a single replicate was enriched for GO term; “ossification” suggesting slight variability of inclusion of surrounding mandible across samples. (Fig. 2A) Marker genes of 5 clusters (4,6,8,12,14) were enriched for epithelial related GO terms including skin development (Fig. 2A). Cells of these clusters also express dental epithelial marker gene *Pitx2* (Mucchielli *et al*., 1997) (Fig. 2B), so were classified “Epithelial Cells” (Fig 2C). Marker genes of 1 cluster are enriched for red blood cell-related GO terms including erythrocyte differentiation (Fig. 2A), strongly express *Hemoglobin 1a* (Fig. 2B), and were thus labelled “Red Blood Cells”. The marker genes of cluster 11 were enriched for GO terms of the vasculature including vasculature development (Fig. 2A). Cells of this cluster express endothelial marker *Cdh5* (Matsumura, Wolff and Petzelbauer, 1997) (Fig. 2B), consistent with the presence of blood vessels in the developing pulp, and were annotated as “Endothelial Cells” (Fig. 2C). The marker genes of cluster 13 displayed enrichment of immune-related GO terms including leukocyte mediated immunity (Fig. 2A). Expression of *Fcer1g* (Fig. 2B) indicates this cluster contains circulating immune cells (Sweet *et al*., 2017), and was classified as “Other Blood Cells” (Fig. 2C). Lastly, marker genes of cluster 15 were enriched for GO terms related to glia, cell migration, and angiogenesis (Fig. 2A, Supp. Table 3). This suggests this small cluster to be perivascular cells (Matsumura, Wolff and Petzelbauer, 1997; Crisan *et al*., 2012) and was annotated “Perivascular Cells” (Fig. 2C).

Overall we identified 6 general cell populations within the developing tooth (Fig. 2C). To validate these classifications we leveraged markers of cell types in the adult mouse incisor (Krivanek *et al*., 2020). We considered published marker genes with at least 2-fold specific enrichment in epithelium, mesenchyme, immune cells, perivascular cells, and endothelium (Supp. Table 4). Marker genes for each cell type were considered as “modules” to calculate the combinatorial activity of groups of genes across our cell populations. Each cluster had the highest correlation with module activity for marker genes of the cognate postnatal incisor cell type (Fig 2D). These results confirm the robustness of our annotations and the persistence of these cell types in the adult tooth.

### Classification of the enamel knot

The identification of the major expected cell types within the developing tooth across multiple biological replicates gave us confidence that we could uncover distinct subpopulations of cells. Specifically, we were interested in the enamel knot due to its potential in tooth regeneration. Its reported small size (Lesot *et al*., 1996, 1999) makes it difficult to isolate manually, but leaves open the possibility of identification through scRNAseq and subsequent *in silico* analyses.

We first interrogated all cell types for expression of genes known to be expressed by the EK. We compiled a list of 98 genes (Supp. Table 5) expressed in the primary mouse enamel knot from biteit (a literature-based dental atlas) and the Jackson Laboratory mouse gene expression database (Smith *et al*., 2019). This list of known EK genes was then used to calculate module scores (KnownEK) for each of the 6 general cell types found in the developing molar (Fig. 2D). We observed significant enrichment of KnownEK scores in the epithelium, consistent with reported epithelial origins of the EK (Fig 3A,B) (Mogollón *et al*., 2021). However, this module is not uniformly enriched across the epithelium, but is concentrated in a subpopulation of the epithelium (Fig 3A).

**Figure 3.**
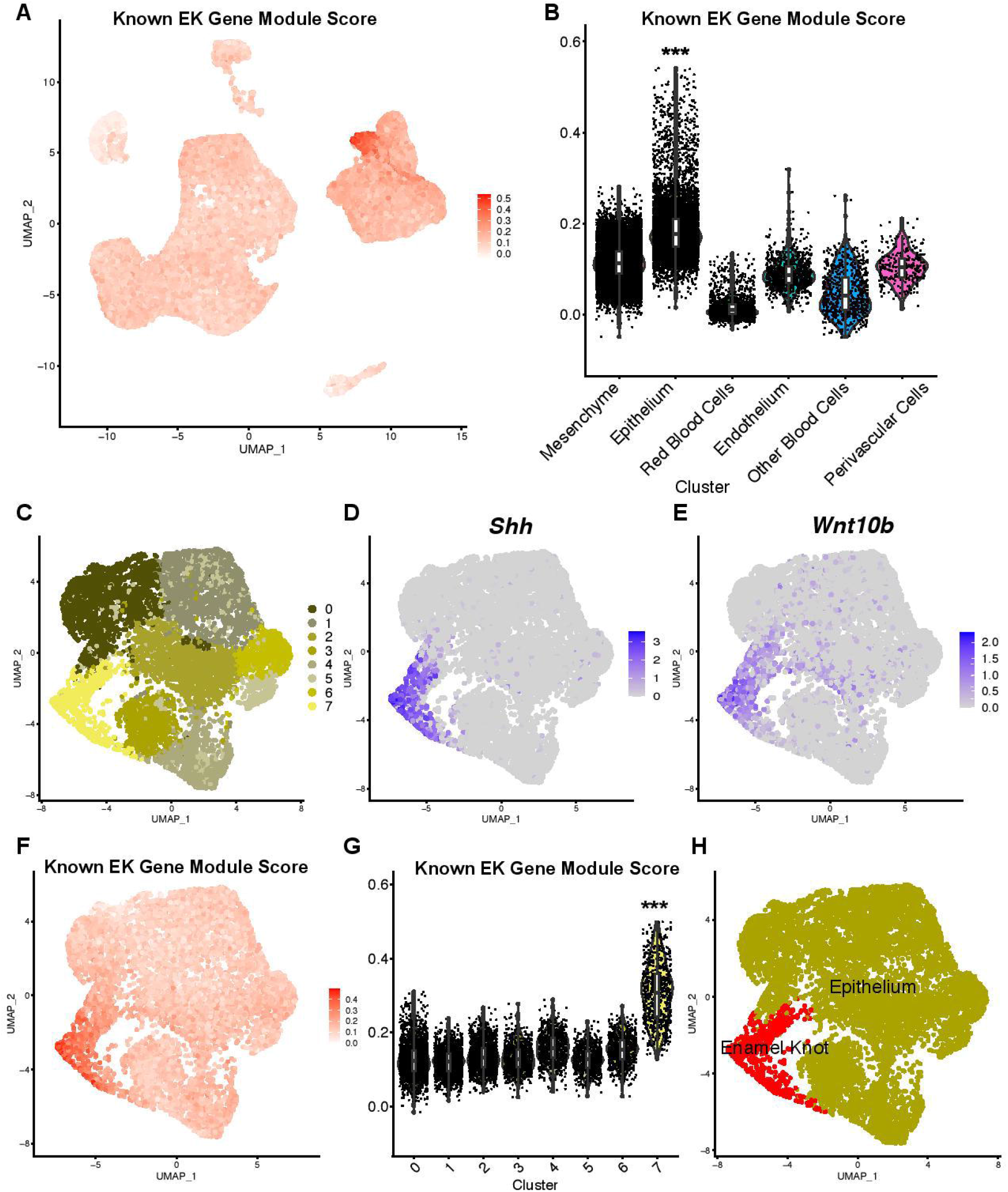
Identification of the Enamel Knot in the Epithelium. **A**. UMAP projection overlaid with KnownEK module score per cell. **B**. Violin plot of A, calculated by annotated cell type shown in Figure 2. P<0.001 for all comparisons using the Wilcoxon test. Boxplot shows median and quartiles. **C.** UMAP projection of epithelial subclustering. **D,E.** Epithelial UMAP projection overlaid with scaled and normalized expression per cell of *Shh* and *Wnt10b*, enamel knot marker genes. **F.** Epithelial UMAP projection of KnownEK module score per cell. **G**. Violin plot of D, calculated by numerical epithelial cluster shown in C. P<0.001 for all comparisons using Wilcoxon test, boxplot shows median and quartiles per group. **H.** Annotated epithelial UMAP projection.

We therefore prioritized our analysis to cells from the epithelial cluster (n=8566). When we considered these cells in isolation and reclustered them, we identified 8 unique epithelial subclusters (Fig. 3C). To initially define cells of the EK, we examined expression of known EK marker genes *Shh* and *Wnt10b* across all epithelial clusters (Vaahtokari *et al*., 1996; Jernvall *et al*., 1998). We observed strong and specific expression in one specific group, cluster 7 (Fig. 3F,G), which we defined as the putative EK cluster (Fig. 3D,E). To confirm this EK classification, we also compared KnownEK module scores across these distinct subpopulations and observed a significant enrichment in cluster 7, consistent with marker gene expression. We also examined cell cycle staging of epithelial clusters, as cells of the EK reportedly express *Cdkn1a* to maintain aproliferative G1 phase despite secreting proliferation-inducing factors (Jernvall *et al*., 1998; Jung *et al*., 2018). We found that 81% of cells (538/661) from the putative EK cluster are maintained in the G1 phase of the cell cycle, further suggesting EK identity. Lastly, we found that the putative EK cluster included an average 7.6% of the total epithelial cells per replicate (SD=0.020), and an average of 1.9% of the total cell count per replicate (SD=0.009, Supp. Fig 4) which generally agrees with calculations based on 3D investigations of the same stage (~5% of cells in the epithelium, Lesot *et al*., 1996, 1999).Taken together, these findings identify this distinct epithelial cluster as cells of the enamel knot; the cluster was re-annotated relative to other epithelial subclusters (Fig. 3H) and relative to other cell types in the developing molar (Fig. 4A).

**Figure 4.**
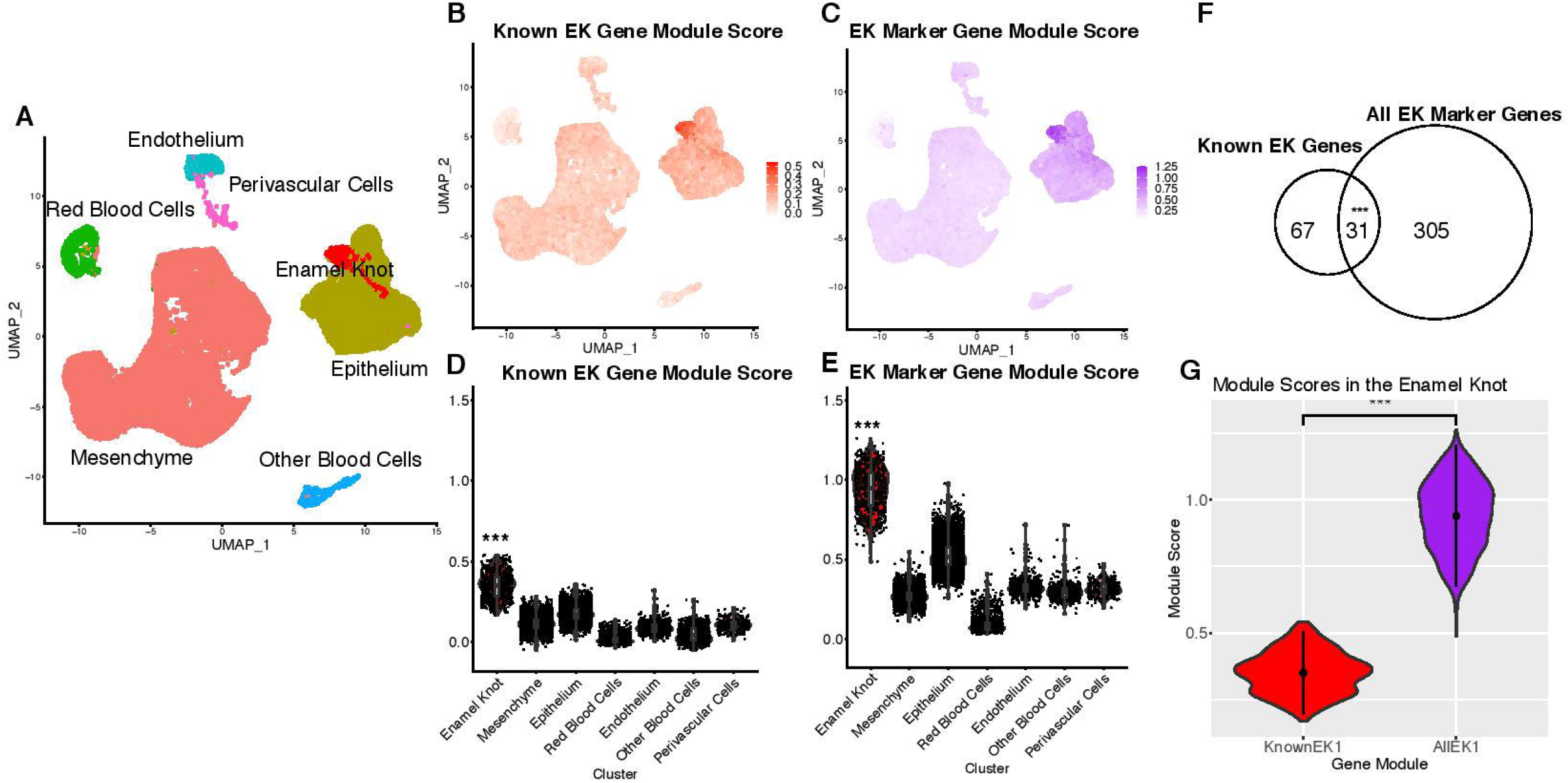
Marker Genes of the Enamel Knot Outperform Known EK Genes. **A**. Annotated UMAP projection featuring cells of the enamel knot identified in Figure 3. **B**. UMAP projection overlaid with KnownEK module score per cell. **C**. Violin plot of B, grouped by annotated cell type including the EK. P<0.001 for all comparisons using Wilcoxon test, boxplot shows median and quartiles. **D**. UMAP projection overlaid with AllEK module score per cell. **E**. Violin plot of D, grouped by annotated cell type including the EK. P<0.001 for all comparisons using Wilcoxon test, boxplot shows median and quartiles. **F.** Euler plot illustration of the overlap between genes known to be expressed in the enamel knot, versus enamel knot marker genes in this investigation. P<0.00001, permutation test. **G.** Direct comparison of module scores of known EK genes and EK marker genes in cells from the enamel knot cluster. P<0.001, Wilcoxon test.

### Identification and Validation of Enamel Knot Marker Genes

While EK cells exhibit high enrichment of KnownEK genes, the expression of this module was observed across multiple cell types (Fig 4B). This is potentially due to EK genes being identified from different stages of development or the qualitative nature of many previous approaches. For instance, the module we assembled included a commonly used canonical EK gene *Bmp4* which is a critical factor in the controlled apoptosis of the EK (Vainio *et al*., 1993; Jernvall *et al*., 1998; Mammoto *et al*., 2011), which is also expressed during some stages in the dental mesenchyme (Supp Fig 3). *Msx2* and *Lef1*, which are also canonical EK genes, showed similar broad expression patterns (Kratochwil *et al*., 1996; Bidder, Latifi and Towler, 1998). We therefore sought to identify a more specific gene signature for the EK and to illuminate potentially novel EK genes. Using a differential expression based approach (Methods), we uncovered 335 EK marker genes with significant enrichment in the putative EK cluster relative to all other clusters (Supp. Table 6). As expected, these markers were enriched for GO terms relevant to tooth development including “odontogenesis” (Supp. Table 7). While EK cells express all KnownEK genes (98/98, p<0.0001; Supp. Fig. 5), only 32% (31/98, p<0.0001; Fig. 4F, Supp. Fig. 5) of KnownEK genes passed our quantitative filters for EK marker gene identification. However, the shared set of 31 genes included well established EK-specific genes *Shh, Cdkn1a*, and *Wnt10b* (Vaahtokari *et al*., 1996; Jernvall *et al*., 1998; Sarkar and Sharpe, 1999). We then considered all EK marker genes from our differential expression analysis as a module (AllEK) and calculated module scores across all clusters. We observed significant enrichment of the AllEK module in the putative EK with significantly higher absolute and relative module scores compared to the KnownEK module (Fig. 4E,F). This indicates that while the KnownEK genes were beneficial for initial identification of the structure, the EK expresses many other structure-specific genes that have not been previously identified.

We then sought to confirm the EK-based expression of the most highly enriched EK marker genes, specifically genes with at least 2 fold enrichment in the structure (64 genes, Top EK Genes, “TEKGs”; Supp. Table 8). To validate these TEKGs, we performed a literature survey and referenced the GenePaint resource of in situ hybridization (ISH) images of sagittal sections of the whole E14 to E14.5 mouse embryo (King *et al*., 1997; Visel, Thaller and Eichele, 2004; Yaylaoglu *et al*., 2005; Diez-Roux *et al*., 2011; Eichele and Diez-Roux, 2011). The 64 TEKGs include 9 known EK genes, including the known EK markers *Shh, Wnt10b*, and *Cdkn1a*. All of these known EK genes showed strong ISH signals in the primordial molar, validating these genes as true markers of the structure.

Of the 55 highly enriched novel EK marker genes, 89% (49/55) had previously been assayed via ISH. We found that the overwhelming majority of these novel genes (42/49) showed strong and specific hybridization in the tooth (Fig. 5, Supp. Table 9), closely mirroring the results for *Shh* expression. Seven genes which were interrogated by ISH showed ubiquitous bodywide expression or available sections at E14.5 did not clearly capture the tooth. An interactive table of all 64 top EK markers, their expression in the tooth, and results of single cell clustering analyses can be visualized and further explored on our laboratory website, https://cotney.research.uchc.edu/scrna-mouse-molar/.

**Figure 5.**
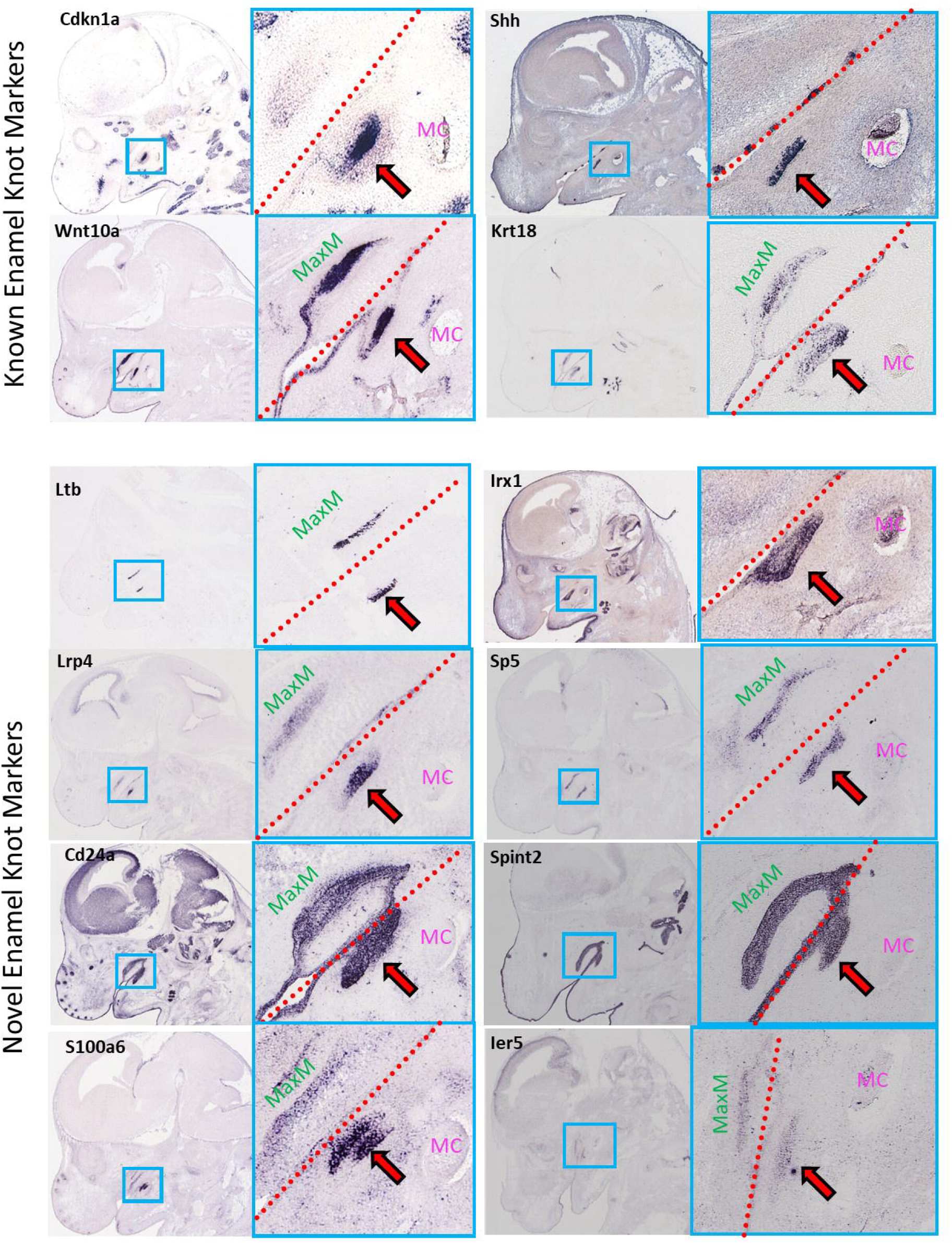
Validation of Novel Highest-Enriched Enamel Knot Markers. *In situ* hybridization images of E14.5 mouse heads courtesy of the GenePaint resource. Blue boxes on sections of the head indicate the approximate area shown in insets. Red dashed line indicates the approximate maxillary/mandibular prominence separation. “MaxM” (green) indicates the maxillary molar, if visible in the section. “MC” indicates Meckel’s cartilage. Red and black arrow indicates the mandibular molar being used for validation. **A-D.** Known enamel knot genes *Cdkn1a, Wnt10a, Shh*, and *Krt18*. **E-L.** Novel enamel knot marker genes *Ltb, Irx1, Lrp4, Sp5, Cd24a, Spint1, S100a6*, and *Ier5*.

## Discussion

Here we have presented our in-depth *in silico* isolation and expression profiling of the enamel knot signalling center in the E14 molar. In this work, we demonstrated that with stringent data processing and rigorous re-analysis pipelines, we can identify small populations of specific cell types within the developing tooth and obtain a gene signature that defines them. Following clustering and systematic marker gene identification, we chose gene ontology based approaches for cell type identification rather than using only a handful of well studied genes. This approach allowed us to functionally annotate populations in a less biased manner than relying on canonical marker genes, which we showed may not be specifically expressed in this assay type. Furthermore, GO-based cell classifications were validated with modules of genes from the adult mouse tooth (Krivanek *et al*., 2020).

With the resulting annotations, we first interrogated cell types for enrichment of a module of genes reported to be expressed or important for EK formation and maintenance. This revealed the strongest enrichment in the epithelial cluster, in agreement with reported EK origins, which allowed us to focus our analysis on this general cell type. We then sub-clustered the epithelium revealing 8 distinct sub-populations (Fig. 3). Given the epithelial nature of all these populations, we did not leverage GO-based cluster annotation; rather, we re-examined the known EK module scores, which were significantly enriched in a single epithelial subcluster. This subcluster showed the highest enrichment of well documented EK specific genes *Shh, Cdkn1a*, and *Wnt10a*, and exhibited enrichment of aproliferative cells, another hallmark of the EK.

Interestingly, while 3 dimensional measurements of the EK have been performed, the absolute and proportional volumes of the molar EK have not been well categorized (Lesot *et al*., 1996, 1999). We found that the size of the putative EK cluster as a proportion of the whole tooth germ (average 1.9%/replicate) generally agreed with 3D-based approximations (<5% of cells)(Lesot *et al*., 1996, 1999). With this evidence, we felt confident annotating these cells as the enamel knot.

While some EK factors are known, a complete transcriptomic profile has not previously been generated. Thus, we identified marker genes whose expression is enriched in and specific to the putative EK cluster compared to all other cell types in the dataset. It is important to consider marker genes in context of all cell types as the EK expresses genes which are contemporaneously expressed in the epithelium and/or mesenchyme (Fig. 4, Supp. Fig. 3), thus excluding them as true EK marker genes. We uncovered 335 EK marker genes, of which only 37 had already been identified as expressed in the EK. These 335 genes were enriched for GO terms consistent with the structure’s epithelial etiology, including “odontogenesis” (Supp. Table 7). When these genes are considered altogether, they show a significantly increased specificity in the EK compared to previously known EK genes (Fig. 4C,F).

Of particular interest are very highly enriched EK markers; 64 genes are enriched >2-fold in the EK, which we called “TEKGs”. The majority of TEKGs (86%; 55/64) did not appear in public databases of known EK genes. We validated the expression of all TEKGs using published literature and hybridization databases. The majority of the TEKGs which had been assayed were seen in the molar at E14.5 with similar patterns to EK marker *Shh* (81%; 52/58; Fig. 5) including all known EK genes (9/9), and the majority of novel TEKGs (86%, 42/49). Our previous work in the developing heart demonstrated an enrichment of disease related genes in those that are most specifically expressed across tissues. (VanOudenhove *et al*., 2020) Therefore we propose these validated, novel TEKGs play an important role in the function of the EK, in tooth development, and potentially tooth-related diseases. Further downstream experiments and analyses will be necessary to unravel their role in tooth patterning and morphology. We provide a convenient portal to explore all of our results at http://cotneyweb.cam.uchc.edu/tooth_scRNA/ with minimal computational effort for the convenience of the research community.

## Methods

Raw fastqs were retrieved from GEO (GSE142201) and aligned to the mm10 genome (GRCm39) using Kallisto/bustools function kbcount (kallisto v0.46.2, bustools v0.40.0)(Melsted *et al*., 2021). Kallisto/bustools was used in filtering and Seurat object generation. Seurat (v3.2.0)(Yaylaoglu *et al*., 2005) was used to merge replicates. Cells were filtered (<5% mtDNA, >200 nFeatureRNA) and log normalized. Cell cycle stages were assigned with mouse orthologs from known human cell cycle genes (Supp. Table 10). Data was scaled using all genes (Supp. Table 2). Nearest neighbors were assigned using dimensions 1:15 via Louvain algorithm. Clusters were generated at resolution=0.3. UMAPs were generated with dimensions 1:15. Epithelial analysis was clustered at resolution=0.2, using dimensions 1:15. Marker genes were identified with DESeq2 using logfc>0.5, min.pct=0.25. SymIDs were converted to EntrezID with clusterProfiler (v3.14.3, org.Mm.eg.db v3.10.0)(Yu *et al*., 2012). clusterProfiler was used to calculate GO term enrichment against background. clusterProfiler “simplify” combined similar GO terms, with a z score cutoff of 0.5. Modules were generated with marker genes logFC>2 from Krivanek et al., 2020 (Supp. Table 4).

Known EK genes were obtained from biteit (*Gene expression in tooth*, 2015) under the heading “Enamel Knot” and from the JAX mouse gene expression database (Smith *et al*., 2019) for genes under the term “Primary Enamel Knot” (EMAPA:38091, filtered by genes with positive assays); gene names were changed to most commonly used gene symbols (i.e. *p21* changed to *Cdkn1a)* and duplicate gene names were merged. In situ hybridization images for EK marker gene validation were obtained from the GenePaint online resource (Visel et al., 2004; Yaylaoglu et al., 2005; Diez-Roux et al., 2001; Eichelle and Diez-Roux, 2011) and a literature search for specific genes in the developing tooth (Nagakawa et al., 2020; King et al., 1997).

Figures were generated using ggplot2 v3.3.3, ggsgnif v0.6.1 and patchwork v1.0.1. ShinyCell (v.2.0.0)(Ouyang *et al*., 2021) package was used for development of the web portal. Scripts for computational analyses are available at https://github.com/cotneylab/scRNA_EnamelKnot.

## Acknowledgments

This work was funded by NIH Grants R03DE028588, R01DE028945, and R35GM119465.

We report no conflicts of interest in this work.

Author 1 contributed to conception, design, data acquisition and interpretation, writing, revising, and creating figures for the manuscript. Author 2 Contributed to conception, design, revising, and creating figures for the manuscript. All authors gave their final approval and agree to be accountable for all aspects of the work.

