## Supplemental Figures for "Identification of enamel knot gene signature within the developing mouse molar"

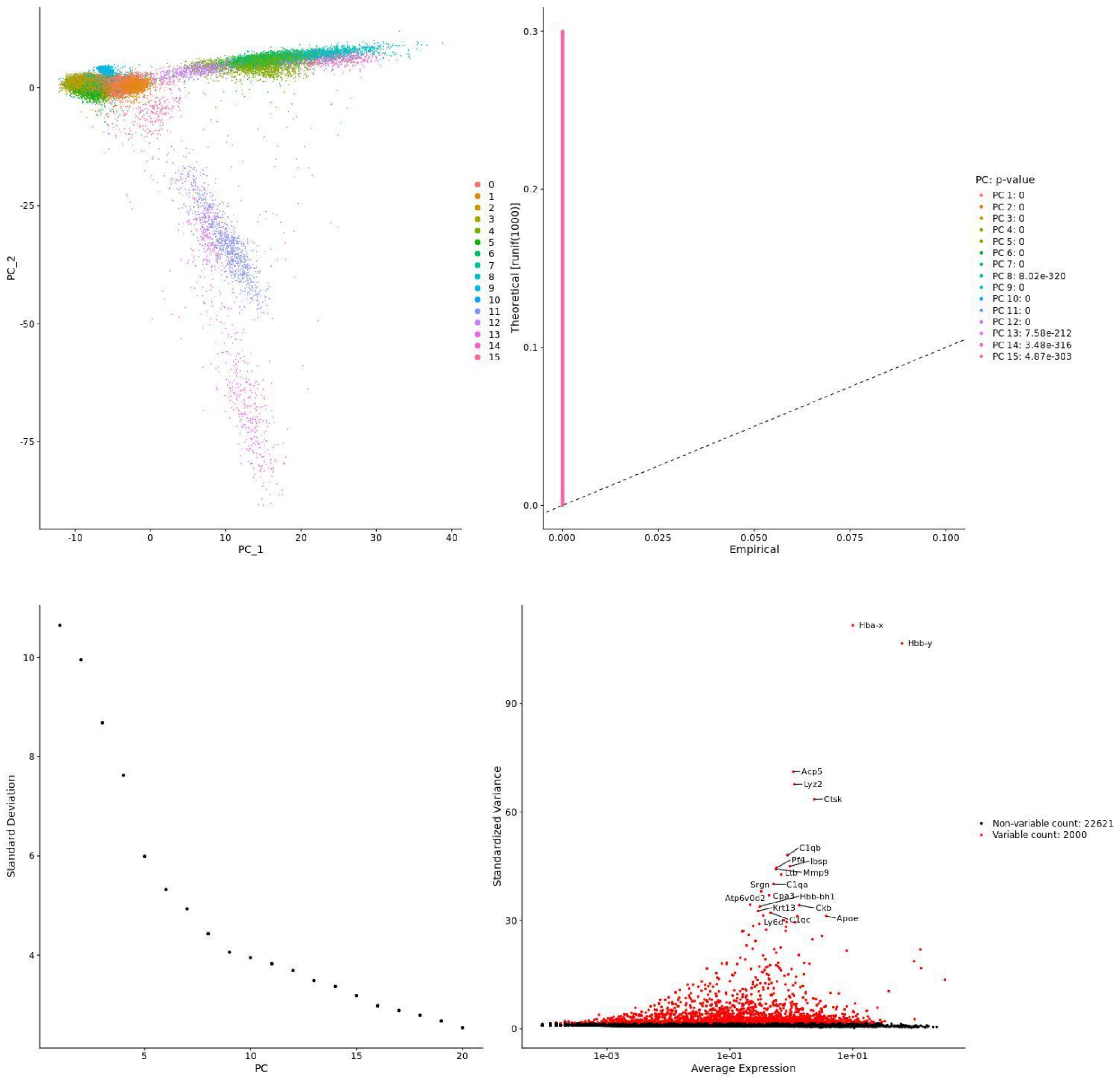


**Supplemental Figure 1. Quality Control Metrics, Dimensionality Calculations, and Variable Feature Identification**

**A.** Principal component analysis (PCA) projection of the first two principal components, cells colored by cluster number. **B.** Jackstraw plot based on dimensions 1:15. **C.** Elbow plot of first 20 dimensions of data. **D.** Featureplot illustrating variability of all genes. Top most variable features labeled.

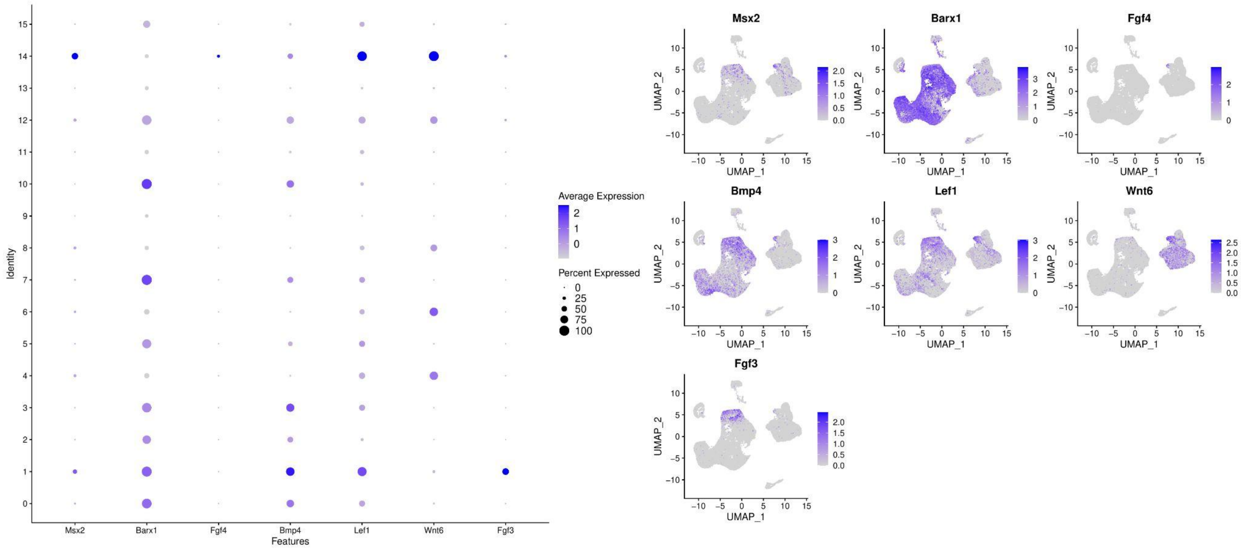


**Supplemental Figure 2. Marker Gene Interrogation of All Clusters**

**A**. Dotplot of numerical cluster annotation vs dental genes of interest. Dot size indicates percent of cells in the cluster expressing the gene, intense blue indicates higher expression level. **B.** UMAP projection overlaid with gene expression scores per cell for genes *Msx2, Barx1, Fgf4, Bmp4, Lef1, Wnt6, Fgf3*.

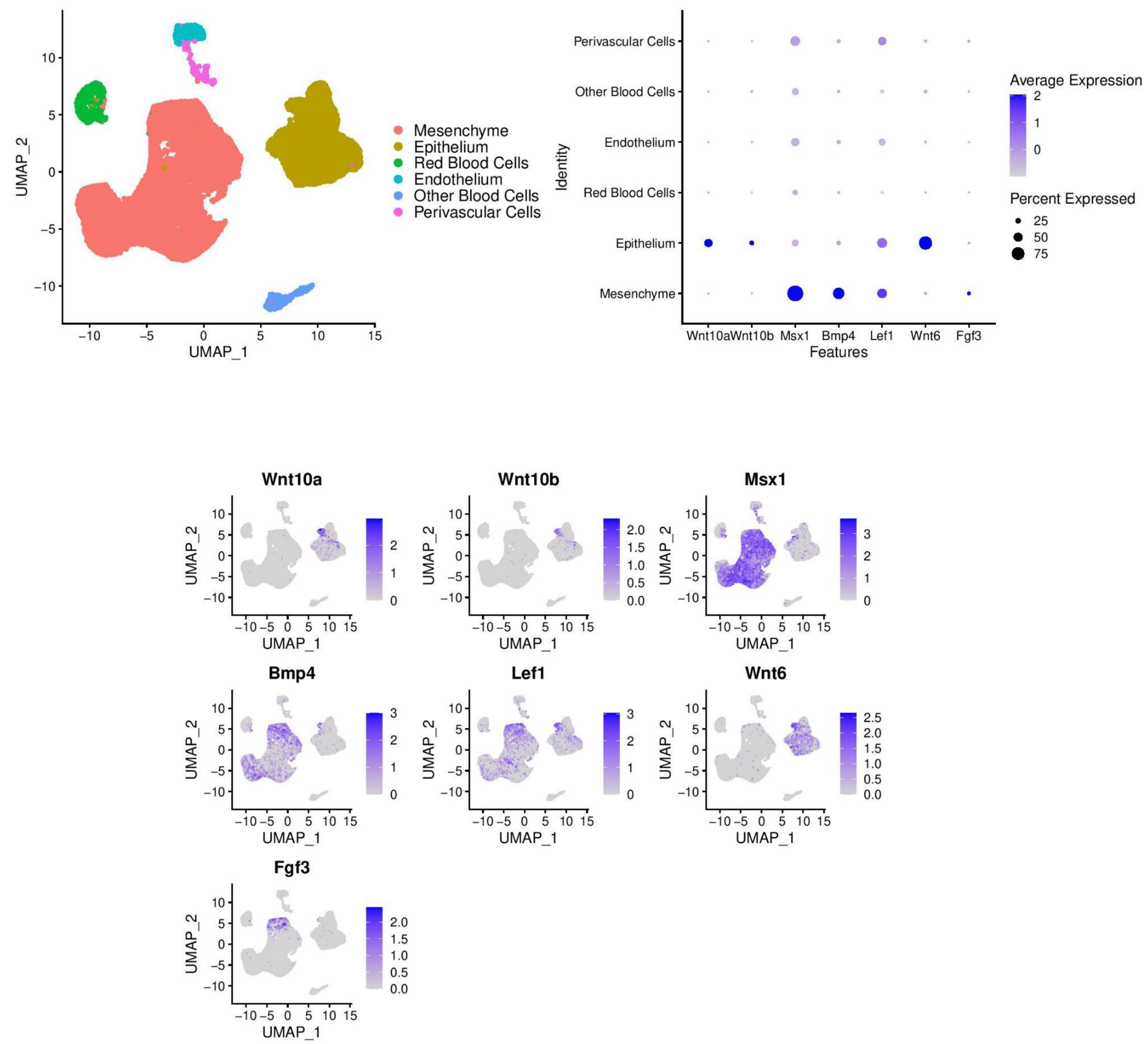


**Supplemental Figure 3. Known Enamel Knot Genes Show Nonspecific Expression in the E14 Molar**

**A**. UMAP projection with overlay of original numerical cluster identities. **B**. Dotplot of annotated cluster versus selected known enamel knot genes. Dot size indicates percent of cells in the cluster expressing the gene, intense blue indicates higher expression level. **C**. UMAP projection overlaid with gene expression scores per cell for known EK genes.

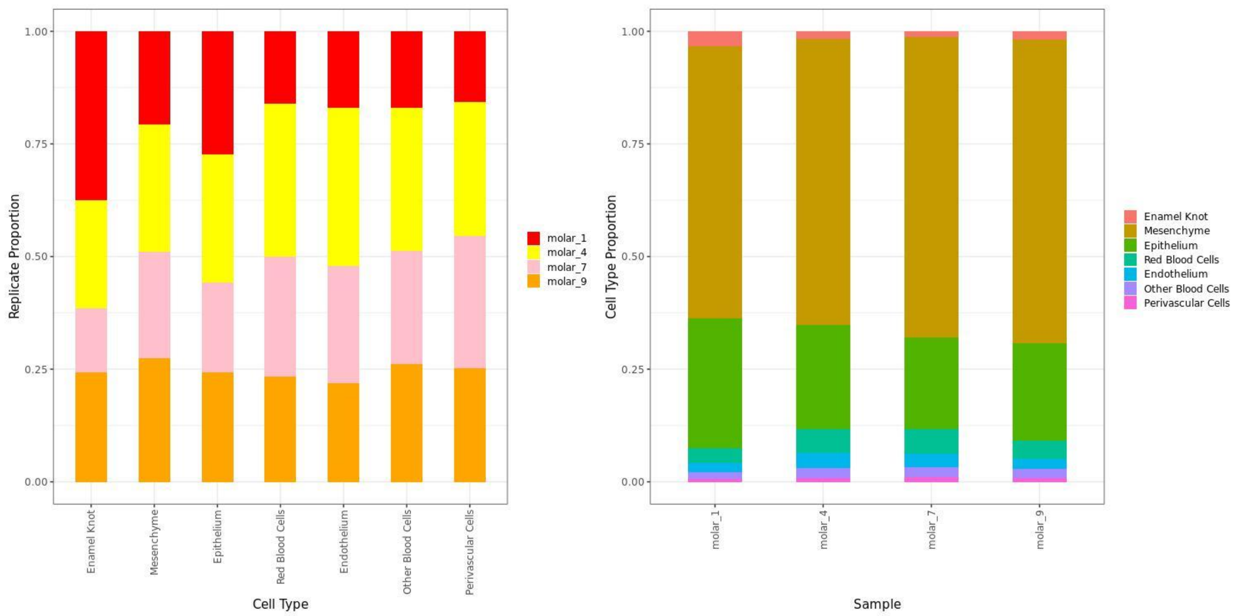


**Supplemental Figure 4. Replicate Contribution to Cells of the Enamel Knot Cluster**
**A.** Per-cluster visualization of replicate contribution to each cell type. **F**. Per-replicate visualization of percent of cells per cell type.

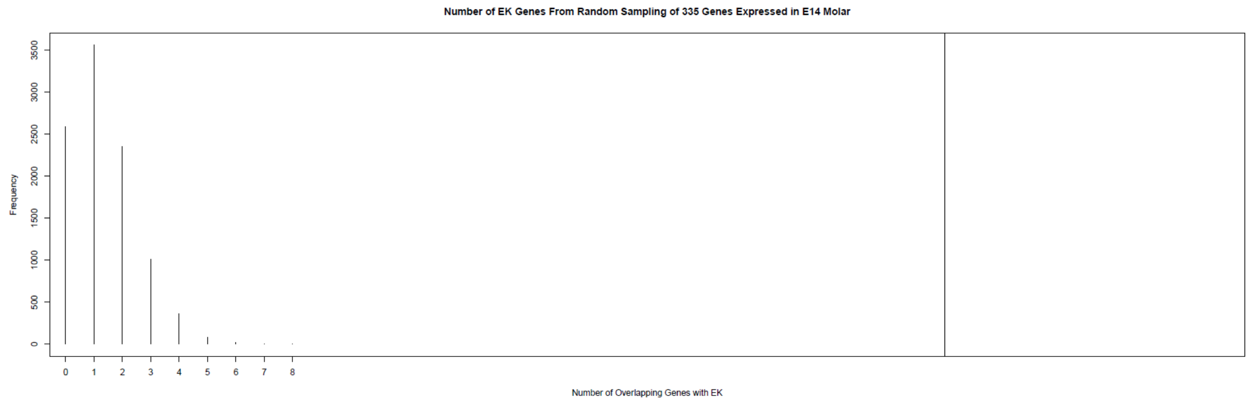


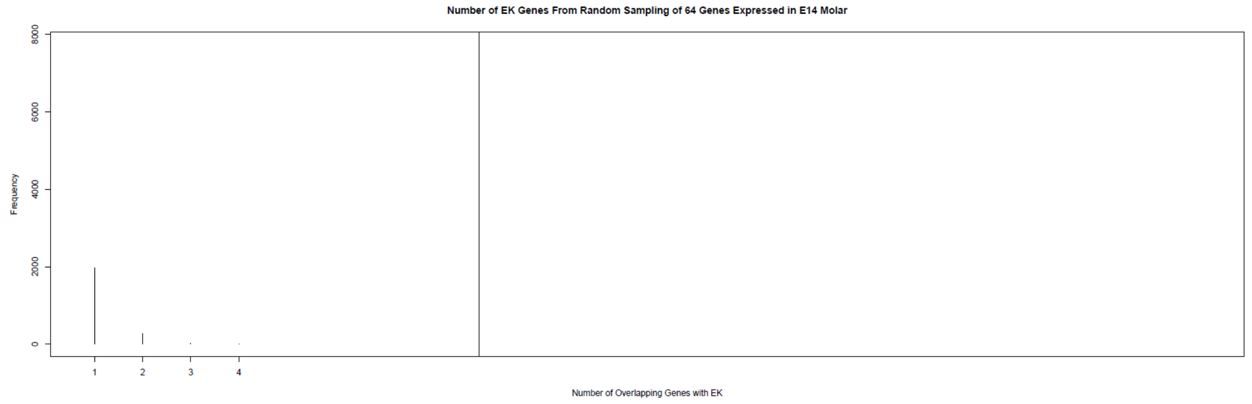

**Supplemental Figure 5. Permutation Analysis of Statistical Significance**

**A.** Permutation analysis (n=10,000) of random sampling of 335 genes from all genes expressed in the E14 molar. 0/10,000 times saw greater than 15 genes on the list of EK marker genes, indicating a $p<0.00001$. **B.** Permutation analysis (n=10,000) of random sampling of 64 genes from all genes expressed in the E14 molar. 0/10,000 times saw greater than 8 genes on the list of top EK marker genes, indicating a $p<0.00001$.
